# Public Health Response to Pan-Resistant *Pseudomonas aeruginosa* Sequence Type 309 with Tandem Guiana Extended Spectrum β-Lactamases – Genetic Characterization and *In Vivo* Efficacy Assessment using Translational Murine Model

**DOI:** 10.1101/2023.11.17.567598

**Authors:** Tyler Lloyd, Christian M. Gill, Jade Aquino Herrera, Dustin Heaton, Munira Shemsu, Kavita K. Trivedi, Andrew Gorzalski, Monica Bender, Mark Pandori, Vici Varghese, David P. Nicolau

**Affiliations:** Alameda County Public Health Department, Oakland, CA; Center for Anti-Infective Research and Development, Hartford Hospital, Hartford, CT; Division of Infectious Diseases, Hartford Hospital, Hartford, CT; Nevada State Public Health Laboratory, Reno, NV; University of California, Berkeley, Berkeley, CA

**Author notes:** **Corresponding Author Information:** David P. Nicolau, PharmD, FCCP, FIDSA, 100 Pearl Street, Hartford, CT 06103. These first authors contributed equally to this article (Tyler Lloyd, Christian M. Gill).

**Keywords:** Antimicrobial, Resistance, Carbapenemase, Pan-Resistant, Whole, Genome, Sequencing, MLST, GES, Translational, Medicine, Murine, Model, Beta-lactamase, ST-309, phylogenetics, SNP, integron, aqcuired

## Abstract

Through phenotypic and genetic profiling, we identified the first pan resistant *Pseudomonas aeruginosa* high-risk sequence type 309 (ST-309) with Guiana Extended Spectrum β-Lactamase (GES) in California, prior to a national outbreak of VIM-GES *Pseudomonas aeruginosa* (1). Our findings highlight detection and use of the translational murine model provide treatment implications for this emerging *Pseudomonas aeruginosa*.

## The Study

Multi-drug resistant *Pseudomonas aeruginosa* is considered a serious threat by the United States Centers for Disease Control. Pan-resistant *Pseudomonas aeruginosa* are uncommon and not well-studied, presenting a challenge to healthcare providers in selecting effective antimicrobial therapies. (4,8,10). The genetics and enzymology of drug resistance increasingly guides treatment decisions in carbapenem-resistant *Pseudomonas*, such as the presence of serine-carbapenemases or metallo-β-lactamases (4). Commercial molecular assays can detect a subset of these genes, but the application of whole genome sequencing provides a priori diagnostic(s) for the identification of rare or unusual resistance genes in highly resistant organisms.

Since June 2017, Alameda County Public Health Department (ACPHD) has mandated reporting of carbapenem-resistant *Klebsiella* spp., *Enterobacter* spp. and *Escherichia coli* to identify these organisms and limit their spread (11). The Alameda County Public Health Laboratory (ACPHL) uses whole genome sequencing to identify major carbapenemase genes in submitted isolates, and together ACPHD’s Communicable Disease (CD) Team and healthcare providers use these data as surveillance tools for antimicrobial resistance tracking to limit transmission opportunities. The laboratory also provides additional sequencing analysis, such as additional resistance gene detection and multi-locus sequence typing. In furtherance of this work, ACPHD also encourages voluntary submission of other carbapenem-resistant isolates for sequencing and analysis.

In January of 2022, ACPHL received a pan-resistant *Pseudomonas aeruginosa* isolated from an immunosuppressed patient’s peritoneal dialysis catheter site. The isolate was initially considered a contaminant, and the patient was discharged. Five days later, the patient was readmitted with abdominal pain. During this admission, the peritoneal dialysis catheter site appeared infected. The patient was treated with vancomycin and ceftolozane/tazobactam, then tapered to oral cephalexin for exit-site cellulitis. Patient developed a fever at 102°F and was restarted on vancomycin and ceftolozane/tazobactam until cultures were negative.

ACPHL’s genetic analysis identified resistance genes to aminoglycosides, cephalosporins, tetracyclines, penicillins, fluoroquinolones, trimethoprim-sulfamethoxazole, and carbapenems. Tandem copies of Guiana Extended Spectrum β-Lactamase (GES) were detected within a class 1 integron, the first such identification of these genes in California (Table 1 and Figure 2). The tandem copies of GES were allelic variants, one copy of GES-19 which has been described to possess typical ESBL activity, and one copy GES-20 which is associated with carbapenemase activity (2,14). We found this isolate belongs to an emerging high-risk sequence type, ST-309, which has been identified in nosocomial outbreaks, but rarely found within the United States (2,6). Initial sequencing analysis was completed within 2 weeks of receiving the isolate, and our data were available 6 weeks before a reference laboratory provided identical results.

**Table 1:**
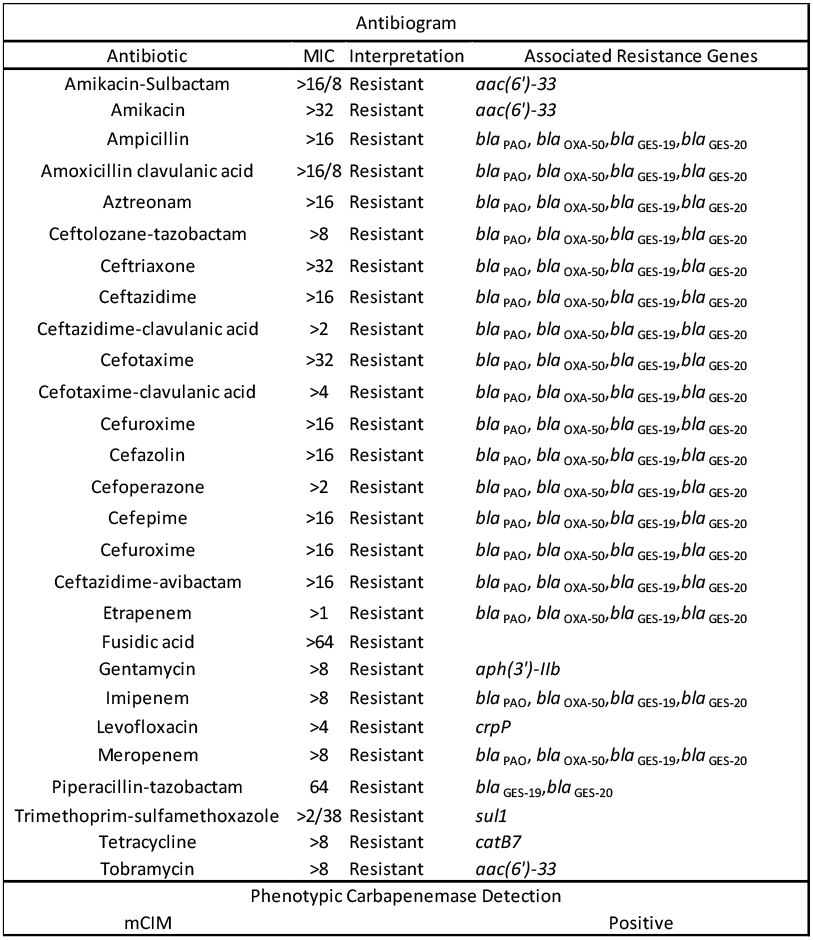
Minimum Inhibitory Concentration (MIC) of Antibiotics Tested *in vitro* of the Guiana Extended Spectrum β-Lactamase, Extensively Drug-Resistant *Pseudomonas aeruginosa* ST-309, Alameda County, California. Drug and drug combinations are listed on the left, MICs are listed in the center in concentrations of µg/mL with CLSI interpretation, and associated resistance genes on the right. MICs from the original clinical report were confirmed by broth microdilution.

In late February of 2022, a second pan-resistant *Pseudomonas aeruginosa* was received at ACPHL. This sample was isolated from urine in a patient with traumatic brain injury who was hospitalized after a fall at a skilled nursing facility. The patient was noted to have urinary retention with blood clots in the Foley catheter on admission. *Pseudomonas aeruginosa* was isolated from a urine sample at this time. Sequencing analysis of this isolate revealed another ST-309 with tandem copies of GES. The CD team was able to identify the isolate as being possibly related, within the same week it was analyzed, to the first case based on the sequence type, resistance profile, temporal proximity to the initial case, and isolate source. Since sequencing data was already available from the first case, ACPHL performed single nucleotide polymorphism (SNP) analysis on request by the CD team to assess for phylogenetic relatedness. We found the initial isolate only differed by 5-6 SNPs from the second isolate, indicating these two isolates were clonally disseminated (Figure 2) (2,15).

Extensively drug-resistant *Pseudomonas aeruginosa* ST-309 has been reported as an emerging high-risk healthcare-associated pathogen. (2,6,9). After case review, no epidemiological links were identified between the cases, despite being closely related. This is important to note because there are no approved *in vitro* diagnostic tests in the United States to detect GES. Therefore, cases of this organism are unlikely to be detected in routine clinical diagnostics. Specific GES alleles also have variable carbapenemase activity, so detection of the gene alone is not sufficient to inform treatment options (14). The ST-309 isolates in this study possess GES alleles with both ESBL and carbapenemase activity. Therapy for hospital-associated *Pseudomonas aeruginosa* infections includes typical anti-pseudomonal β-lactams (e.g., cefepime and meropenem) or the novel β-lactam/ β-lactamase-inhibitor combinations (e.g., ceftolozane/tazobactam and ceftazidime/avibactam) (7). Detection of the high-risk clones described in this report is alarming as the tandem GES-genotype detected conferred resistance to all these agents. Since limited clinical data are available to guide treatment decisions if such isolates are detected in the clinic, translational *in vivo* models provide empiric data to support rational therapeutic choices.

The Center for Anti-Infective Research and Development (CAIRD) at Hartford Hospital has developed and validated a translational *in vivo* murine thigh model with human-simulated exposures of conventionally employed antibiotic regimens to assess their clinical utility in the management of difficult-to-treat pathogens like *Pseudomonas aeruginosa*.(3) When challenged with our clinical isolates, our results demonstrated that a clinically utilized ceftazidime/avibactam alone regimen had variable degrees of *in vivo* killing against the isolates (−3.48±0.27,-1.26±1.66 log_10_ CFU/Thigh), and that a high-dose meropenem regimen given alone showed no activity as defined by growth similar to untreated controls (2.62±0.19, 2.43±0.58 log_10_ CFU/Thigh). The addition of meropenem with ceftazidime/avibactam enhances effectiveness profoundly (−3.27±0.25, -2.96±0.00 log_10_ CFU/Thigh). Cefiderocol monotherapy also showed strong *in vivo* efficacy (−3.36±0.25, -3.07±0.00 log_10_ CFU/Thigh), and was the only drug in which *in vitro* potency was predictive of efficacy (Figure 1).

**Figure 1:**
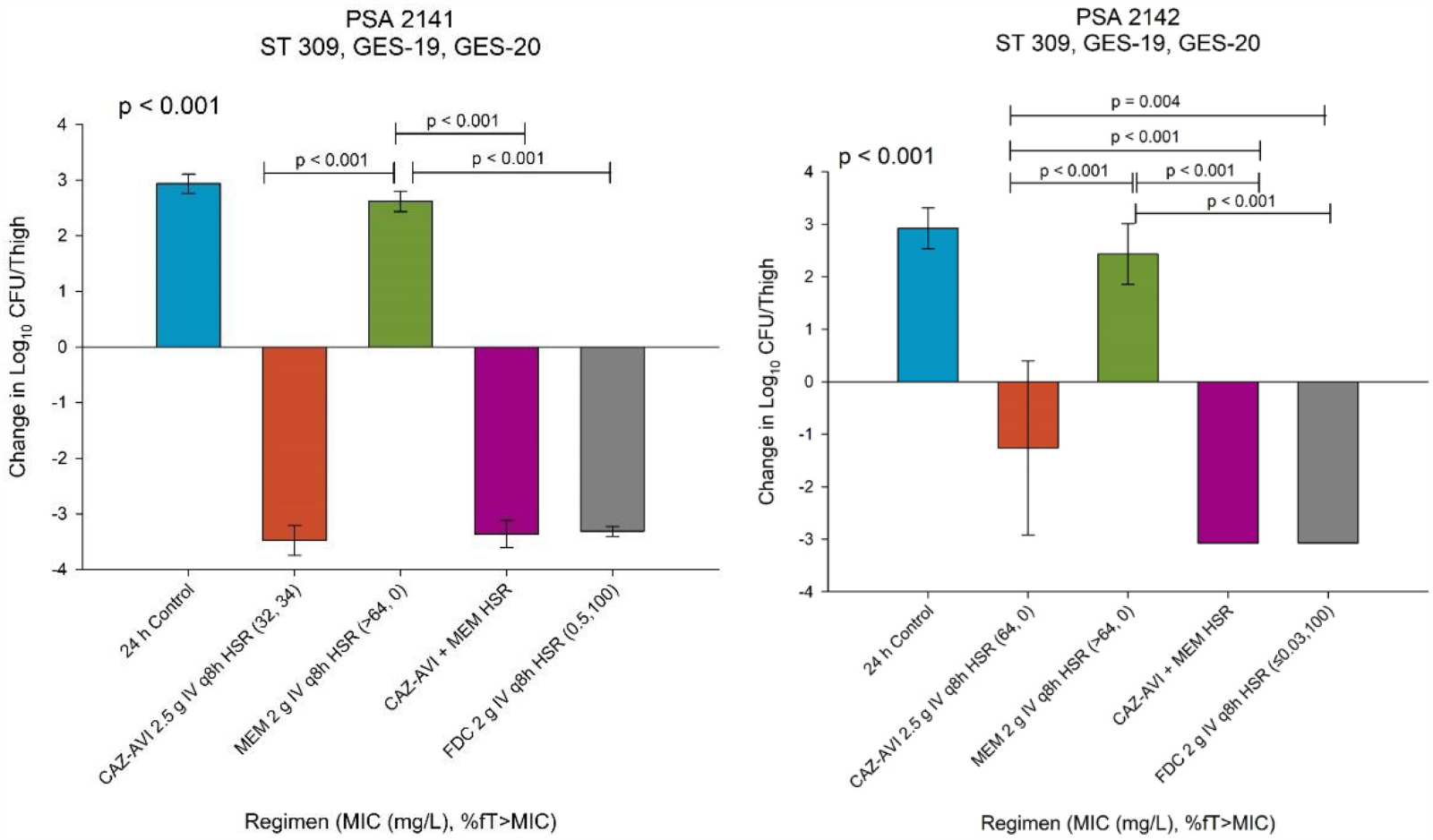
*In vivo* efficacy studies of human-simulated regimens (HSR) of conventionally utilized regimens of Ceftazidime/Avibactam 2.5g Q8 2hr infusion, Meropenem 2g q8 3 hr infusion, combination Ceftazidime/Avibactam + Meropenem, and Cefiderocol 2g q8 3 hr infusion. Results for the 24-hour control and each treatment group represent Change in log_10_ CFU/Thigh compared with baseline bacterial burden (0 h control). Achievement of ≥ 1-log_10_ reduction in bacterial density from the baseline burden in this translational model is associated with clinical success in man. %*f*T>MIC is the duration of free drug concentration above the MIC of the target pathogen.

**Figure 2:**
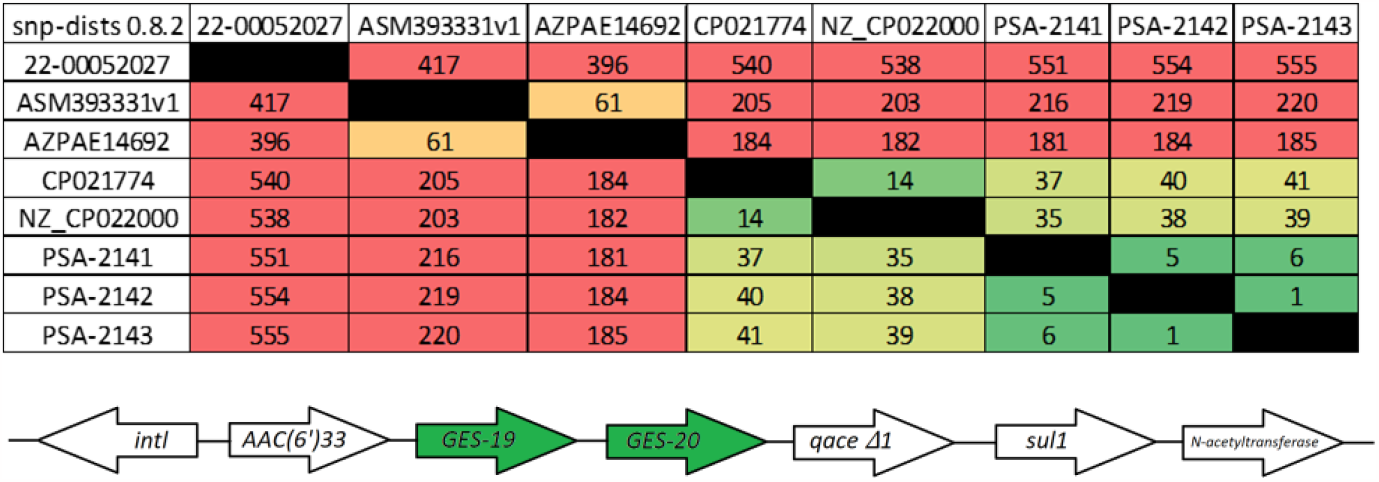
SNP matrix and structure of class one integron housing aminoglycoside resistance (*AAC(6’) 33*), tandem copies of *GES*, quaternary ammonia resistance (*qace Δ1*), and trimethoprim-sulfamethoxazole resistance (*sul1*). PSA-2142 and PSA-2143 represent duplicate samples of the second case.

## Conclusions

Antibiotic stewardship and antimicrobial susceptibility surveillance tools remain core strategies against our ability to identify, track and prevent transmission of drug-resistant bacteria. Our results demonstrate that an inherent quality of routine whole genome sequencing is the ability to identify highly resistant isolates of interest in research for drug therapies. Performing this work in a local public health laboratory can reduce the time to coordinate response efforts and research on effective treatments for pan-resistant *Pseudomonas aeruginosa*. Sequencing as a diagnostic tool also presents an opportunity for building institutional relationships into more effective networks for fighting the spread of antimicrobial resistance. We demonstrated through *in vivo* experiments that ceftazidime/avibactam with meropenem is an effective therapy for pan-resistant ST-309 *Pseudomonas aeruginosa* with tandem GES and reaffirm the efficacy of cefiderocol both *in vitro* and *in vivo* (Figure 1). This study expands the scientific knowledge for appropriate therapy choice for pan-resistant ST-309 *Pseudomonas aeruginosa* and highlights the necessity for surveillance of carbapenem-resistant *P. aeruginosa* for genotypic evidence of GES as such enzymes are likely under described in the United States.

## Biosketch

Tyler Lloyd is a public health microbiologist with the Alameda County Department of Public Health and a PhD student at the University of California, Berkeley. His primary research interests include antimicrobial resistance and bacterial evolution through a public health lens.

